# A murine model with JAK2V617F expression in both hematopoietic cells and vascular endothelial cells recapitulates the key features of human myeloproliferative neoplasm

**DOI:** 10.1101/2021.08.24.457585

**Authors:** Haotian Zhang, Amar Yeware, Sandy Lee, Huichun Zhan

**Affiliations:** Graduate Program in Molecular & Cellular Biology, Stony Brook University, Stony Brook, NY; Department of Medicine, Stony Brook School of Medicine, Stony Brook, NY; Graduate Program in Molecular & Cellular Pharmacology, Stony Brook University, Stony Brook, NY; Medical Service, Northport VA Medical Center, Northport, NY

## Abstract

The myeloproliferative neoplasms (MPNs) are characterized by an expansion of the neoplastic hematopoietic stem/progenitor cells (HSPC) and an increased risk of cardiovascular complications. The acquired kinase mutation JAK2V617F is present in hematopoietic cells in a majority of patients with MPNs. Vascular endothelial cells (ECs) carrying the JAK2V617F mutation can also be detected in patients with MPNs. In this study, we show that a murine model with both JAK2V617F-bearing hematopoietic cells and JAK2V617F-bearing vascular ECs recapitulated all the key features of the human MPN disease, which include disease transformation from essential thrombocythemia to myelofibrosis, extramedullary splenic hematopoiesis, and spontaneous cardiovascular complications. During aging and MPN disease progression, there was a loss of both HSPC number and HSPC function in the marrow while the neoplastic hematopoiesis was relatively maintained in the spleen, mimicking the advanced phases of human MPN disease. Different vascular niche of the marrow and spleen could contribute to the different JAK2V617F mutant stem cell functions we have observed in this JAK2V617F-positive murine model. Compared to other MPN murine models reported so far, our studies demonstrate that endothelial dysfunction plays an important role in both the hematologic and cardiovascular abnormalities of MPN.

**Key Points:**

- A murine model in which JAK2V617F is expressed in both hematopoietic cells and ECs recapitulated the key features of the human MPN disease
- Different vascular niche of the marrow and spleen could contribute to different JAK2V617F HSC functions during MPN disease progression

## Introduction

The myeloproliferative neoplasms (MPNs), which include polycythemia vera (PV), essential thrombocythemia (ET), and primary myelofibrosis (PMF), are clonal stem cell disorders characterized by hematopoietic stem/progenitor cell (HSPC) expansion, overproduction of mature blood cells, a tendency to extramedullary hematopoiesis, an increased risk of transformation to acute leukemia or myelofibrosis, and an increased risk of vascular thrombosis^1,2^. The incidence of MPNs increases significantly with aging and MPN is uncommon before the age of 50 years^3^. Older age and longer disease duration are also associated with higher risk of disease transformation to myelofibrosis or secondary acute myeloid leukemia, as well as increased morbidity and mortality in these patients^4^. These observations suggest that aging plays an important role in MPN development.

The acquired signaling kinase mutation JAK2V617F is present in most patients with MPNs and aberrant JAK-STAT signaling plays a central role in these disorders^5^. Although JAK2V617F-positive murine models have provided unequivocal evidence that JAK2V617F is able to cause MPNs, there is significant heterogeneity in disease phenotypes between different murine models, and none has been able to recapitulate both the myeloproliferative phenotype and the cardiovascular pathology in patients with MPNs^6^. In addition, these murine models were mostly followed for less than 3-9 months^7–18^ and how aging affects MPN disease progression has not been studied.

Endothelial cells (ECs) are an essential component of the hematopoietic niche and most HSPCs reside close to a marrow sinusoid (the “perivascular niche”)^19^. Vascular ECs also play critical roles in the regulation of hemostasis and thrombosis^20^. The JAK2V617F mutation can be detected in microvascular ECs isolated from liver and spleen (by laser microdissection), and marrow (by flow cytometry sorting) in 60-70% of patients with MPNs^21–23^. The mutation can also be detected in 60-80% of EC progenitors derived from the hematopoietic lineage and, in some reports based on *in vitro* culture assays, in endothelial colony-forming cells from patients with MPNs^22–26^. Previously, we reported that the JAK2V617F-bearing vascular endothelium promotes the expansion of the JAK2V617F mutant HSPCs in preference to wild-type HSPCs^27–31^ and contributes to the development of cardiovascular complications^32^ in a murine model of MPN. In the present study, we investigated how MPN progresses in the JAK2V617F-bearing vascular niche during aging.

## Materials and Methods

### Experimental mice

JAK2V617F Flip-Flop (FF1) mice^12^ was provided by Radek Skoda (University Hospital, Basal, Switzerland) and *Tie2-Cre* mice^33^ by Mark Ginsberg (University of California, San Diego). FF1 mice were crossed with Tie2-Cre mice to express JAK2V617F specifically in all hematopoietic cells (including HSPCs) and vascular ECs (Tie2^+/-^ FF1^+/-^, or Tie2FF1), so as to model the human diseases in which both the hematopoietic stem cells and ECs harbor the mutation. All mice used were crossed onto a C57BL/6 background and bred in a pathogen-free mouse facility at Stony Brook University. Animal experiments were performed in accordance with the guidelines provided by the Institutional Animal Care and Use Committee.

### Marrow and spleen cell isolation

Murine femurs and tibias were first harvested and cleaned thoroughly. Marrow cells were flushed into PBS with 2% fetal bovine serum using a 25G needle and syringe. Remaining bones were crushed with a mortar and pestle followed by enzymatic digestion with DNase I (25U/ml) and Collagenase D (1mg/ml) at 37 °C for 20 min under gentle rocking. Tissue suspensions were thoroughly homogenized by gentle and repeated mixing using 10ml pipette to facilitate dissociation of cellular aggregates. Resulting cell suspensions were then filtered through a 40uM cell strainer.

Murine spleens were collected and placed into a 40uM cell strainer. The plunger end of a 1ml syringe was used to mash the spleen through the cell strainer into a collecting dish. 5ml PBS with 2% FBS was used to rinse the cell strainer and the resulting spleen cell suspension was passed through a 5ml syringe with a 23G needle several times to further eliminate small cell clumps.

### Complete blood counts and in vitro assays

Complete blood counts and hematopoietic colony formation assays were performed as we previously described^34^. Mouse methylcellulose complete media (Stem Cell Technologies, Vancouver, BC) was used to assay hematopoietic colony formation, which was enumerated according to the manufacturer’s protocol.

For Lineage negative (Lin^-^) cell culture, marrow or splenic Lin^-^ cells were first enriched using the Lineage Cell Depletion Kit (Miltenyi Biotec). On Day 0, 1,000 Lin^-^ cells were seeded in a 48-well plate and cultured in 150ul StemSpan serum-free expansion medium (SFEM) containing recombinant mouse Stem cell factor (100 ng/ml), recombinant mouse Interleukin-3 (6 ng/ml) and recombinant human Interleukin-6 (10 ng/ml) (all from Stem Cell Technologies). 200ul fresh SFEM medium with cytokines was added on Day 5 and 8 and cells were counted on Day 5 and 10.

### Histology

Femur and spleen tissues were fixed in cold 4% paraformaldehyde for 6hr at 4°C while shaking. The tissues were washed with PBS for 8-16hrs at room temperature to remove paraformaldehyde. Femurs were then decalcified and paraffin sections (5-μm thickness) were stained with hematoxylin and eosin or reticulin (Reticulum II Staining Kit, Roche, Tucson, AZ) to assess fibrosis. Images were taken using a Nikon Eclipse TS2R inverted microscope (Nikon, Melville, NY).

### Flow cytometry

All samples were analyzed by flow cytometry using a FACSAriaTM III or a LSR II (BD biosciences, San Jose, CA, USA). Lineage cocktail (include CD3, B220, Gr1, CD11b, Ter119; Biolegend), cKit (Clone 2B8, Biolegend), Sca1 (Clone D7, Biolegend), CD150 (Clone mShad150, eBioscience), CD48 (Clone HM48-1, Biolegend), CD45 (Clone 104) (Biolegend, San Diego, CA, USA), and CD31 (Clone 390, BD biosciences) antibodies were used.

### BrdU incorporation analysis

Mice were injected intraperitoneally with a single dose of 5-bromo-2’-deoxyuridine (BrdU; 100 mg/kg body weight) and maintained on 1mg BrdU/ml drinking water for two days. Mice were then euthanized and marrow cells isolated as described above. For analysis of HSC (Lin^-^cKit^+^Sca1^+^CD150^+^CD48^-^) proliferation, Lin^-^ cells were first enriched using the Lineage Cell Depletion Kit (Miltenyi Biotec) before staining with fluorescent antibodies specific for cell surface HSC markers, followed by fixation and permeabilization using the Cytofix/Cytoperm kit (BD Biosciences, San Jose, CA), DNase digestion (Sigma, St. Louis, MO), and anti-BrdU antibody (Biolegend, San Diego, CA) staining to analyze BrdU incorporation^31^.

### Analysis of apoptosis by active caspase-3 staining

Marrow cells were stained with fluorescent antibodies specific for cell surface HSC markers, followed by fixation and permeabilization using the Cytofix/Cytoperm kit (BD Biosciences). Cells were then stained using a rabbit anti-activated caspase-3 antibody^31^. Data were acquired using a LSR II flow cytometer.

### Analysis of senescence by senescence associated β-galactosidase (SA-β-Gal) activity

Marrow cells were stained with fluorescent antibodies specific for cell surface HSC markers. Cells were then washed and fixed using 2% paraformaldehyde and incubated with CellEvent^™^ Senescence Green Probe (ThermoFisher Scientific, Waltham, MA) according to the manufacturer’s instruction. Data were acquired using a LSR II flow cytometer.

### VE-cadherin in vivo staining and immunofluorescence imaging

25ug Alexa Fluor 647-conjugated monoclonal antibodies that target mouse VE-cadherin (clone BV13, Biolegend) were injected retro-orbitally into 2yr old Pf4^+^FF1^+^ or control mice under anesthesia^35^. Ten minutes after antibody injection, the mice were euthanized. Mouse femurs and spleens were dissected out and washed in PBS. After fixation in 4% paraformaldehyde (PFA) (Affymetrix) for 6hr at 4°C while rotating, the samples were washed in PBS overnight to remove PFA, cryoprotected in 20% sucrose, embedded in OCT compound (Tissue-Tek), and flash frozen at -80°C. Frozen samples were cryosectioned (~10uM) using a Leica CM1510S Cryostat. Images were acquired using a Nikon Eclipse Ts2R inverted fluorescence microscope.

### Transthoracic echocardiography

Transthoracic echocardiography was performed on mildly anesthetized spontaneously breathing mice (sedated by inhalation of 1% isoflurane, 1 L/min oxygen), using a Vevo 3100 high-resolution imaging system (VisualSonics Inc., Toronto, Canada). Both parasternal long-axis and sequential parasternal short-axis views were obtained to assess global and regional wall motion. Left ventricular (LV) dimensions at end-systole and end-diastole and fractional shortening (percent change in LV diameter normalized to end-diastole) were measured from the parasternal long-axis view using linear measurements of the LV at the level of the mitral leaflet tips during diastole. LV ejection fraction (EF), volume, and mass are measured and calculated using standard formulas for the evaluation of LV systolic function^32,36^.

### Histology

Hearts and lungs were fixed in 4% PFA overnight at 4°C while rotating. The tissues were then washed multiple times with PBS at room temperature to remove PFA. Paraffin sections (5-μm thickness) were stained with Hematoxylin/Eosin (H&E) following standard protocols. Images were taken using a Nikon Eclipse Ts2R inverted microscope.

### Statistical Analysis

Statistical analyses were performed using Student’s unpaired, 2-tailed *t* tests using Excel software (Microsoft). A *p* value of less than 0.05 was considered significant. For all bar graphs, data are presented as mean ± standard error of the mean (SEM).

## Results

### The Tie2FF1 mice develop ET to PMF transformation during aging

The Tie2FF1 mice developed an ET-like phenotype with neutrophilia (3.8 vs 1.8 × 10^3^/uL, *P*=0.014), thrombocytosis (1068 vs 558 × 10^3^/uL, *P*=0.036), and normal hemoglobin at 2mo of age, results consistent with previous reports^28,37^. We followed these mice up to 18mo of age to evaluate how the JAK2V617F mutant vascular niche regulate MPN neoplastic hematopoiesis and disease transformation during aging. The Tie2FF1 mice continued to develop increasing neutrophilia and thrombocytosis, both of which plateaued at ~1 yr of age. In addition, the mice developed significant lymphocytosis and anemia after 6mo of age (Figure 1A-B). At 18mo of age, there was significant splenomegaly (spleen weight 611mg vs 89mg, *P*<0.001), increased total spleen cell counts (267 vs 153 × 10^6^ cells per spleen, *P*=0.031), and decreased total marrow cell counts (23 vs 58 × 10^6^ cells per femur, *P*<0.001) in the Tie2FF1 mice compared to age-matched Tie2-cre control mice (Figure 1C-E). Histology examination revealed extensive marrow osteopetrosis and destroyed splenic architecture, as well as increased fibrosis in both the marrow and spleen of the old Tie2FF1 mice compared to age matched control mice (Figure 1F-I). No evidence of leukemia transformation was observed in the Tie2FF1 mice. These findings indicate that the Tie2FF1 mice developed ET to PMF disease transformation with extramedullary splenic hematopoiesis during aging.

**Figure 1.**
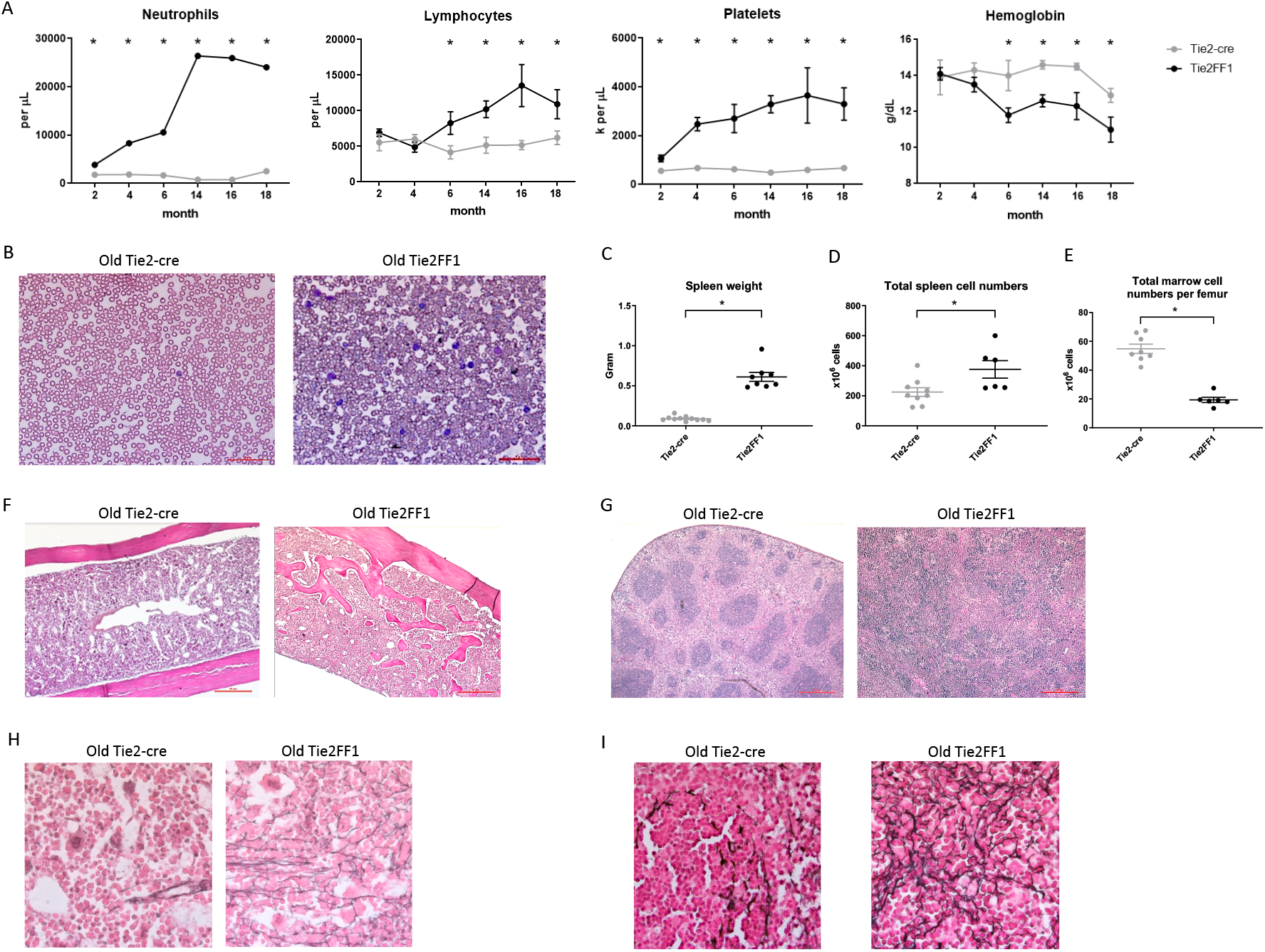
The Tie2FF1 mice develop ET to PMF transformation during aging. (**A**) Peripheral blood cell counts of Tie2FF1 (black line) and Tie2-cre control mice (grey line). (n=5-10 mice in each group at 2mo and 4mo; n=6-7 mice in each group at 6mo, 12mo, and 15mo; n=10 mice in each group at 18mo). (**B**) Representative peripheral blood smear of 18mo old Tie2-cre and Tie2FF1 mice (40X magnification). (**C-E**) Spleen weight (C), total spleen cell counts (D), and total femur cell counts (E) in 18mo old Tie2-cre and Tie2FF1 mice (n=10 mice in each group). (**F-G**) Representative hematoxylin and eosin sections of marrow (F) and spleen (G) from 18mo old Tie2-cre and Tie2FF1 mice (F: 10X magnification; G: 4X magnification). (**H-I**) Representative reticulin stain of marrow (H) and spleen (I) from 18mo old Tie2-cre and Tie2FF1 mice (40X magnification). * *P* <0.05

### Decreased marrow hematopoiesis during aging in the Tie2FF1 mice

Previously, we and others reported that marrow HSCs were significantly expanded in young Tie2FF1 mice compared to age-matched Tie2-cre control mice^28,31,38^. To examine how aging affects the neoplastic hematopoiesis in MPN, we first measured the numbers of marrow hematopoietic progenitor cells using colony formation assays. We found that the total hematopoietic progenitor cells were significantly increased in young (4-5mo) Tie2FF1 mice compared to age-matched control mice (2.6-fold, *P*=0.005); in contrast, there was no significant difference in progenitor cell numbers between old (18mo) Tie2FF1 mice and age-matched control mice (Figure 2A). Consistent with these findings, flow cytometry analysis revealed that marrow Lin^-^cKit^+^Sca1^+^CD150^+^CD48^-^ HSCs^39^ were significantly expanded in young Tie2FF1 mice compared to age-matched control mice (9.7-fold, *P*<0.001), while there was no significant difference in HSC frequency between old Tie2FF1 and control mice (0.125% vs. 0.184%, *P*=0.087) (Figure 2B). Considering that the total marrow cells were decreased 2.5-fold in the old Tie2FF1 mice (Figure 1E), there was a 3.7-fold decrease in the absolute marrow HSC cell numbers in old Tie2FF1 mice compared to age-matched control mice.

**Figure 2.**
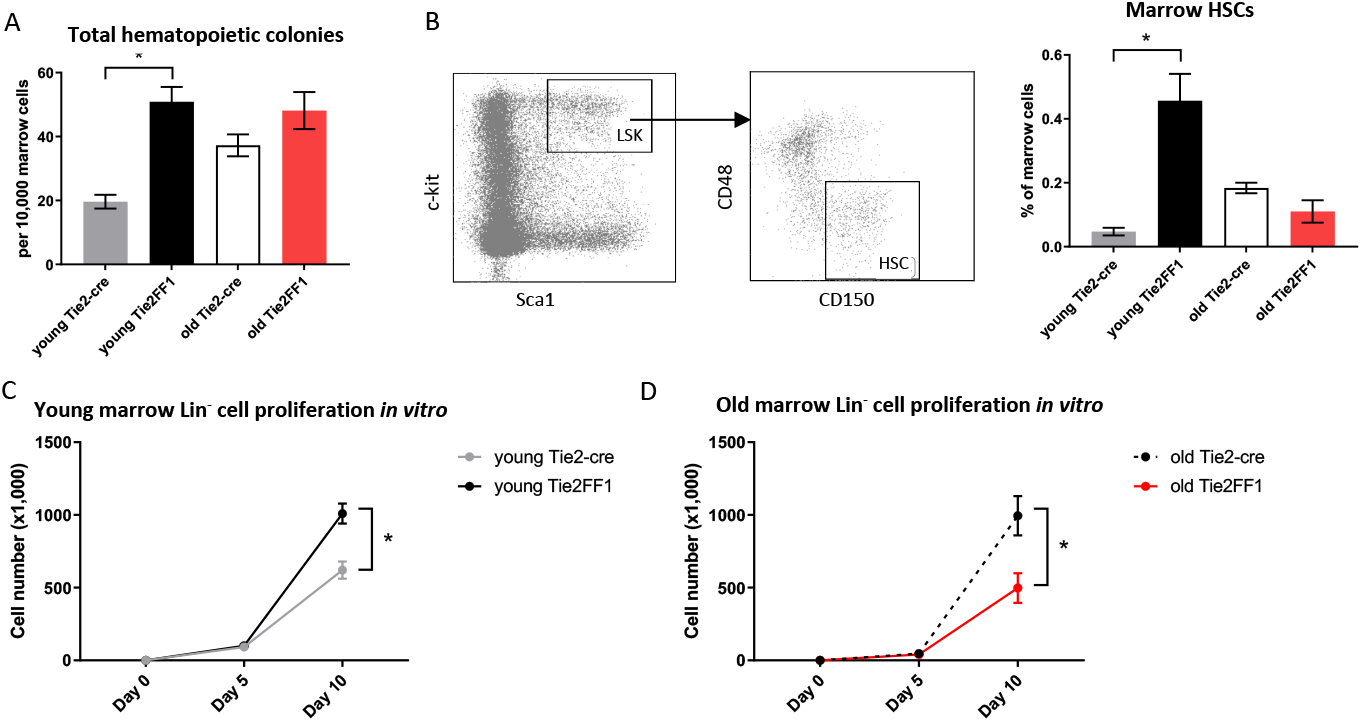
Decreased marrow hematopoiesis in the Tie2FF1 mice during aging. (**A**) Colony formation assays in marrow cells isolated from young (n=4 mice in each group) and old (n=6-7 mice in each group) Tie2-cre control and Tie2FF1 mice. (**B**) Representative flow cytometry plots showing gating strategy (left) of marrow Lin^-^cKit^+^Sca1^+^CD150^+^CD48^-^ HSCs frequency (right) in young (n=7 mice in each group) and old (n=5-6 mice in each group) Tie2-cre control and Tie2FF1 mice. (**C-D**) Cell proliferation of marrow Lin-HSPCs isolated from young (C) and old (D) Tie2-cre control and Tie2FF1 mice. Cells were cultured on SFEM medium containing recombinant mouse SCF (100ng/mL), recombinant mouse IL3 (6ng/mL), and recombinant human IL6 (10ng/mL). Data are from one of two independent experiments (with triplicates in each experiment) that gave similar results. * *P*<0.05

To test whether the decreased phenotypic HSC number was associated with altered HSC function, we isolated marrow Lin^-^ HSPCs from Tie2-cre control and Tie2FF1 mice and measured their cell proliferation *in vitro* in serum-free liquid medium. At the end of a ten-day culture, while JAK2V617F mutant HSPCs from young Tie2FF1 mice displayed a higher proliferation rate than wild-type HSPCs from young control mice (1.6-fold, *P*=0.012), mutant HSPCs from old Tie2FF1 mice proliferated less than old control HSPCs (2.0-fold, *P*=0.043) (Figure 2C-D). Taken together, these data indicated that there was a loss of both HSPC number and HSPC function in the marrow of old Tie2FF1 mice during aging, mimicking the advanced phases of myelofibrosis^3^.

### Expanded splenic extramedullary hematopoiesis in the Tie2FF1 mice

Spleen is the most frequent organ involved in extramedullary hematopoiesis in patients with MPNs^3^. To examine how the splenic hematopoiesis changes during aging and MPN disease progression in the old Tie2FF1 mice, we first measured the numbers of splenic hematopoietic progenitor cells using colony formation assays. We found that splenic hematopoietic progenitor cells were markedly increased in both young (19-fold, *P*<0.001) and old (19-fold, *P*<0.001) Tie2FF1 mice compared to age-matched control mice (Figure 3A). In line with this finding, flow cytometry analysis revealed that spleen HSCs were expanded in both young (53-fold *P* = 0.046) and old (6.2-fold, *P* = 0.004) Tie2FF1 mice compared to age-matched control mice (Figure 3B). Considering that the total spleen cells were increased 1.7-fold in old Tie2FF1 mice (Figure 1D), there was a 10.5-fold increase in the absolute spleen HSC numbers in old Tie2FF1 mice compared to age-matched control mice. When we isolated spleen Lin^-^ HSPCs from young and old Tie2-cre control and Tie2FF1 mice and cultured them *in vitro*, we found that both young (12.9-fold, *P*=0.005) and old (4.3-fold, *P*=0.022) JAK2V617F mutant spleen HSPCs displayed a higher proliferation rate than age-matched wild-type control spleen HSPCs (Figure 3C-D). These results suggest that, in contrast to the decreased HSPC number and HSPC function we have observed in the marrow (Figure 2), the spleen of old Tie2FF1 mice was able to maintain the expansion of JAK2V617F mutant hematopoiesis during aging and MPN disease progression.

**Figure 3.**
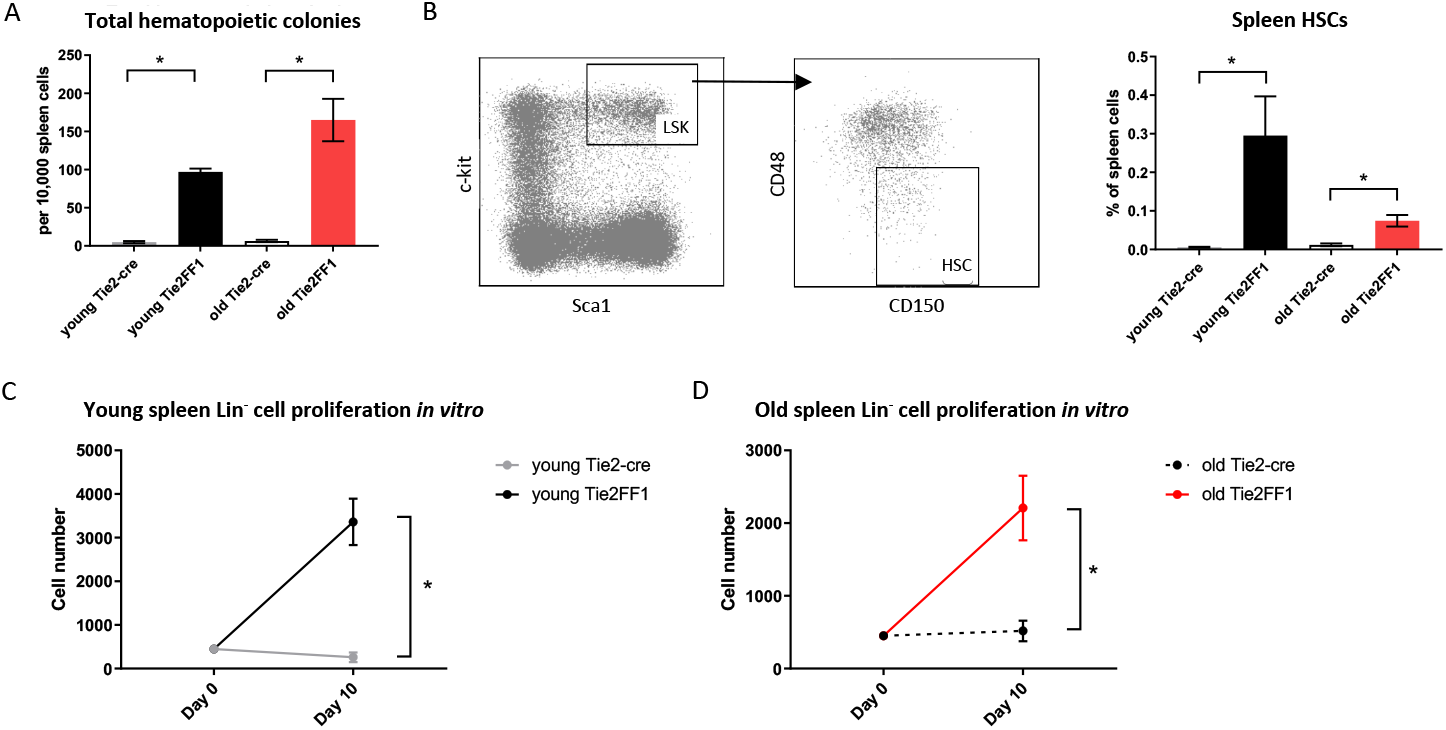
Expanded splenic extramedullary hematopoiesis in the Tie2FF1 mice. (**A**) Colony formation assays in spleen cells isolated from young (n=3 mice in each group) and old (n=5-6 mice in each group) Tie2-cre control and Tie2FF1 mice. (**B**) Spleen Lin^-^cKit^+^Sca1^+^CD150^+^CD48^-^ HSCs frequency in young (n=3 mice in each group) and old (n=5 mice in each group) Tie2-cre control and Tie2FF1 mice. (**C-D**) Cell proliferation of spleen Lin-HSPCs isolated from young (C) and old (D) Tie2-cre control and Tie2FF1 mice. Cells were cultured in SFEM medium containing recombinant mouse SCF (100ng/mL), recombinant mouse IL3 (6ng/mL), and recombinant human IL6 (10ng/mL). Data are from one of two independent experiments (with triplicates in each experiment) that gave similar results. * *P*<0.05

### Different HSC functions in the marrow and spleen of old Tie2FF1 mice

The differences between marrow (Figure 2) and spleen (Figure 3) hematopoiesis in the old Tie2FF1 mice prompted us to further investigate how aging and MPN disease progression affect the JAK2V617F mutant HSC function differently in the marrow and spleen. First, we measured HSC proliferation *in vivo* by BrdU labeling^31^. We found that JAK2V617F mutant HSCs from old Tie2FF1 mice proliferated more rapidly than wild-type HSCs from age-matched control mice in both the marrow (58% vs 21%, *P*=0.001) and the spleen (41% vs 22%, *P*=0.042) (Figure 4A-B). Next, we measured HSC cell apoptosis by assessing their activated caspase-3 levels using flow cytometry analysis^31^. We found that JAK2V617F mutant marrow HSCs from old Tie2FF1 mice displayed higher level of apoptosis compared to wild-type marrow HSCs from age-matched control mice (3.0% vs 0.8%, *P*=0.006); in contrast, mutant spleen HSCs from old Tie2FF1 mice displayed significantly less apoptosis compared to wild-type spleen HSCs from control mice (1.8% vs 11.6%, *P*=0.002) (Figure 4C-D). Since oncogenic mutation is a major stress to induce cellular senescence^40^ and the JAK-STAT signaling has been reported to induce cellular senescence^41–44^, we also measured HSC senescence by measuring senescence associated β-galactosidase (SA-β-Gal) activity using flow cytometry analysis. JAK2V617F mutant marrow HSCs from old Tie2FF1 mice demonstrated significantly higher senescence rates compared to wild-type marrow HSCs from age-matched control mice (18% vs 9%, *P*=0.011); in contrast, there was no difference in the cellular senescence rate between the mutant spleen HSCs from old Tie2FF1 mice and wild-type spleen HSCs from control mice (Figure 4E-F). Taken together, although the JAK2V617F mutant HSCs from old Tie2FF1 mice were more proliferative than wild-type HSCs in both the marrow and spleen, mutant HSCs were more apoptotic and senescent than wild-type HSCs in the marrow while the mutant cells were relatively protected in the spleen.

**Figure 4.**
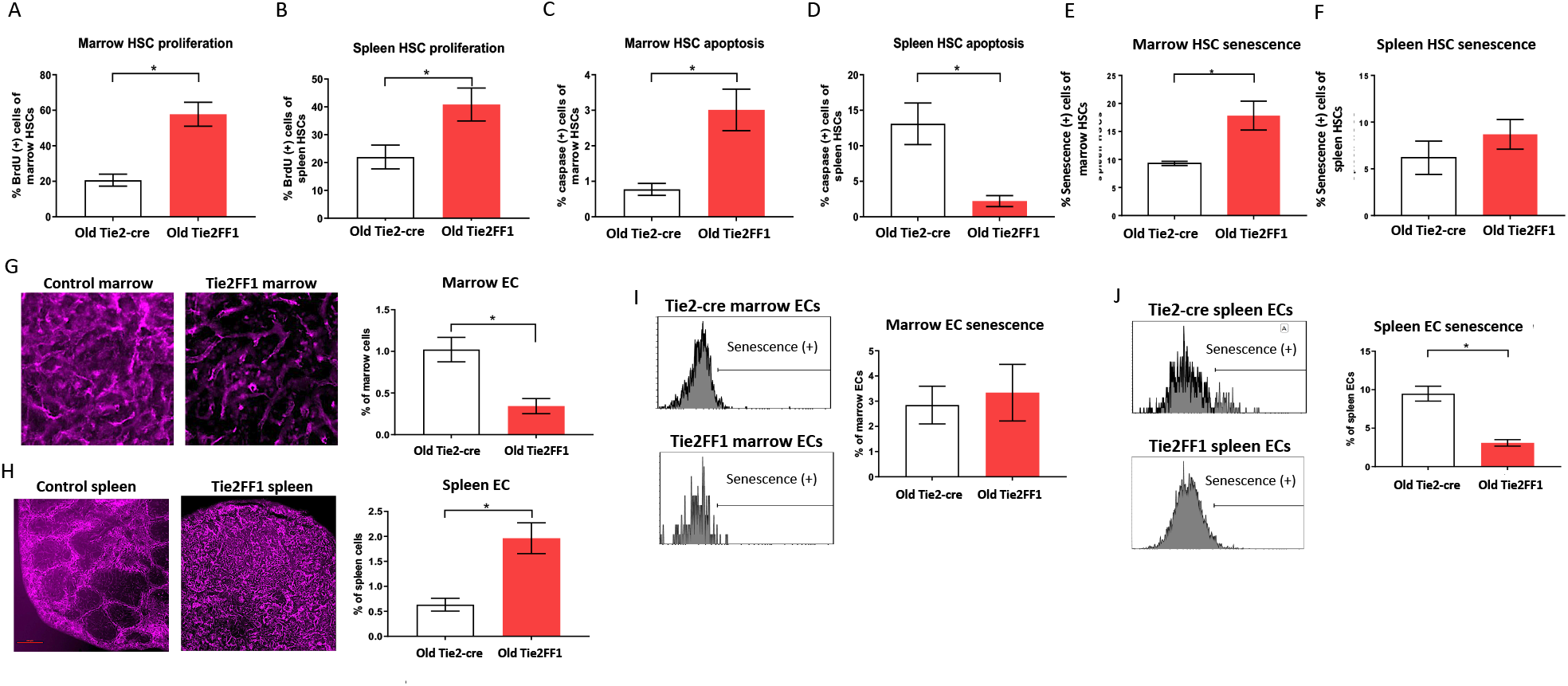
Different HSC functions in the marrow and spleen of old Tie2FF1 mice (**A-B**) Cell proliferation rate of HSCs in the marrow (A) and spleen (B) measured by *in vivo* BrdU labeling (A: n=4-5 mice in each group; B: n=4 mice in each group). (**C-D**) Cellular apoptosis rate of HSCs in the marrow (C) and spleen (D) measured by activated caspase-3 staining using flow cytometry analysis (C: n=5 mice in each group; D: n=5-6 mice in each group). (**E-F**) Cellular senescence rate of HSCs in the marrow (E) and spleen (F) measured by SA-β-Gal activity using flow cytometry analysis (E: n=5 mice in each group; F: n=5-6 mice in each group). (**G-H**) Representative immunofluorescent images of VE-cadherin (+) vasculatures (magnification 10x for femur marrow and 4x for spleen) (left) and flow cytometry quantitative analysis of CD45^-^CD31^+^ ECs (right) of the marrow (G) and spleen (H) from 18mo Tie2-cre control and Tie2FF1 mice (G: n=8 mice in each group; H: n=5 mice in each group). (**I**) Representative flow cytometry histograms (left) and quantitative analysis of cellular senescence of marrow ECs (n=4 mice in each group). (**J**) Representative flow cytometry histograms (left) and quantitative analysis of cellular senescence of spleen ECs (n=3 mice in each group). * *P* < 0.05

Most HSCs reside close to a perivascular niche in the marrow and spleen^19,45^. To understand how different vascular niches contribute to different HSC functions in the old Tie2FF1 mice, we measured marrow and spleen ECs (CD45^-^CD31^+^) by flow cytometry analysis. We found that marrow ECs were significantly decreased in old Tie2FF1 mice compared to age-matched control mice; in contrast, spleen ECs were significantly expanded in old Tie2FF1 mice. These results were also confirmed by *in vivo* VE-cadherin labeling and immunofluorescence imaging of the marrow and spleen tissue samples (Figure 4G-H). In addition, while there was no difference in marrow EC senescence rate between old Tie2FF1 mice and old control mice, the JAK2V617F mutant splenic ECs from old Tie2FF1 mice were much less senescent compared to wild-type splenic ECs from age-matched control mice (Figure 4I-J). Therefore, the different vascular niche of the marrow and spleen could contribute to the decreased marrow hematopoiesis and expanded splenic hematopoiesis we have observed in the Tie2FF1 mice during aging.

### Persistent but compensated cardiomyopathy in the old Tie2FF1 mice

Cardiovascular complications are the leading cause of morbidity and mortality in patients with MPNs. Previously, we reported that the Tie2FF1 mice developed spontaneous heart failure with thrombosis, vasculopathy, and cardiomyopathy at 20wk of age^32^. Here, we followed the cardiovascular function of Tie2FF1 mice during aging. At 18mo of age, the Tie2FF1 mice continued to demonstrate a phenotype of dilated cardiomyopathy with a moderate but significant decrease in LV EF (56% versus 66%, *P*=0.043), an increase in LV end-diastolic volume (84μL vs 71μL, *P*=0.086) and end-systolic volume (38 μL vs 25 μL, *P*=0.041), and an increase in LV mass (156mg vs 113mg, *P*=0.021) compared to age-matched control mice (Figure 5A). Pathological evaluation confirmed the diagnosis of dilated cardiomyopathy in old Tie2FF1 mice with significantly increased heart weight-to-tibia length ratio compared to age-matched control mice (0.014 vs 0.011 gram/mm, *P*=0.004) (Figure 5B). Increased lung weight in old Tie2FF1 mice compared to control mice (0.287 vs 0.226 gram) further indicated the presence of pulmonary edema commonly associated with heart failure (Figure 5C). Similar to what we previously reported in the young Tie2FF1 mice^32^, there was spontaneous thrombosis in the right ventricle and pulmonary arteries in the old Tie2FF1 mice, while age-matched Tie2-cre control mice had no evidence of spontaneous thrombosis in their heart or lungs (Figure 5D-E). Despite these cardiovascular dysfunctions, there was no difference in body weight between old Tie2FF1 mice and control mice (Figure 5F), nor was there any significantly increased incidence of sudden death in the old Tie2FF1 mice compared to age-matched control mice. These findings suggested that there was a persistent but compensated cardiomyopathy and heart failure in the Tie2FF1 mice during aging.

**Figure 5.**
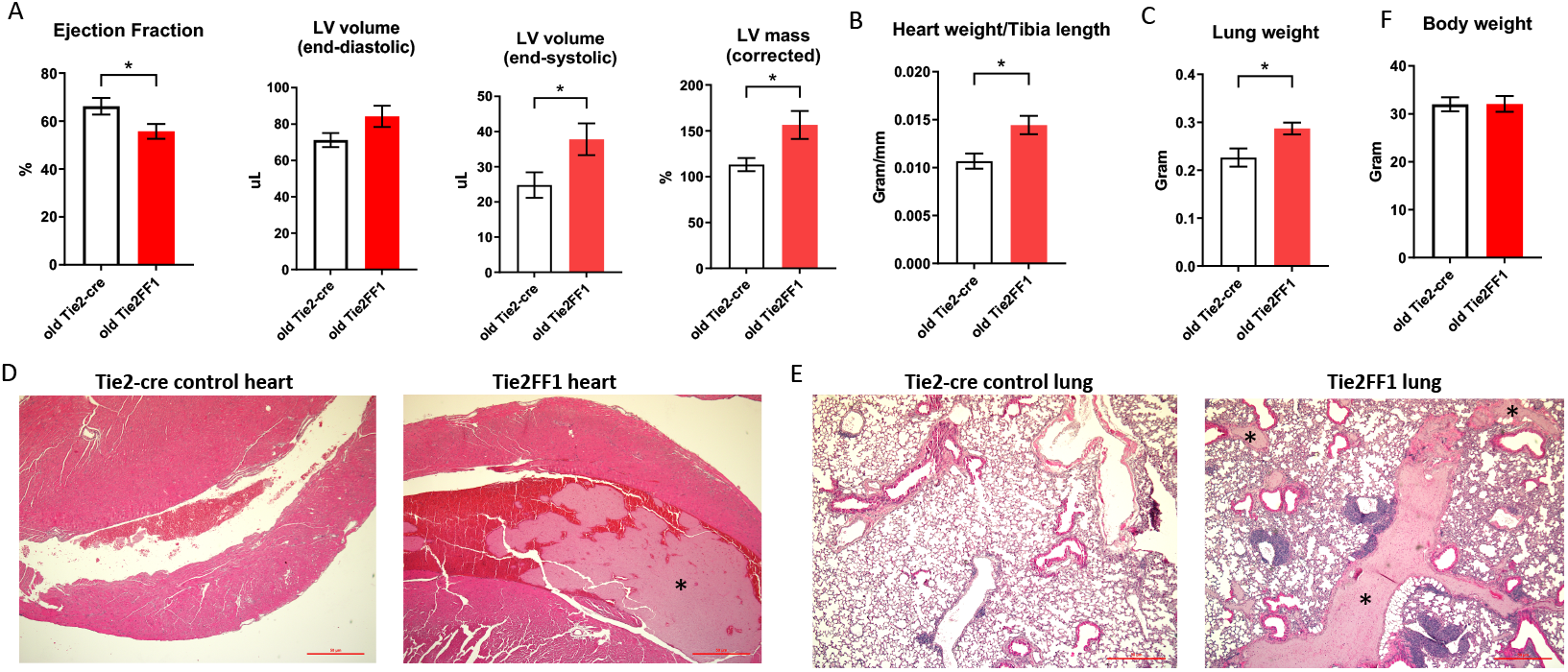
Persistent cardiomyopathy and congestive heart failure in the old Tie2FF1 mice. (**A**) Measurements of left ventricular (LV) ejection fraction, end-diastolic and end-systolic volume, and mass by transthoracic echocardiography in 18mo old Tie2-cre control and Tie2FF1 mice. (**B**) Heart weight adjusted by tibia length of 18mo old Tie2-cre control and Tie2FF1 mice. (**C**) Lung weight of 18mo old Tie2-cre control and Tie2FF1 mice. (**D**) Representative H&E staining of transverse sections of heart from Tie2-cre control and Tie2FF1 mice. Note the presence of thrombus (star *) in right ventricle (magnification 10x). (**E**) Representative H&E staining of coronal sections of lung from Tie2-cre control and Tie2FF1 mice. Note the presence of thrombus (star *) in segment pulmonary arteries of the Tie2FF1 mice (magnification 10x). (**F**) Body weight of 18mo old Tie2-cre control and Tie2FF1 mice. n=8-11 mice in each group. * *P* < 0.05

## Discussions

With heterozygous human JAK2V617F transgene expression in both the hematopoietic cells and vascular ECs, the Tie2FF1 mice developed an ET-like phenotype at young age (2mo old) which transformed to PMF during aging. The mice also demonstrated features of extramedullary splenic hematopoiesis, spontaneous vascular thrombosis, cardiovascular dysfunction, which persisted during the aging process. Compared to other MPN murine models reported so far^7–18^, the Tie2FF1 mice is the first MPN murine model that faithfully recapitulated almost all the key features of the human MPN diseases. Considering the presence of the JAK2V617F mutation in microvascular ECs isolated from patients with MPNs^21–23^ and the recapitulation of all the key features of human MPN diseases by the Tie2FF1 mice, the roles of endothelial dysfunction in the hematologic and cardiovascular pathogenesis of MPN and whether the MPN vascular niche can be targeted to provide more effective therapeutic strategies for patients with these diseases shall be further investigated.

Extramedullary splenic hematopoiesis often compensates for normal hematopoietic suppression in neoplastic conditions^46,47^. Splenomegaly is a common feature in patients with MPNs as a result of extramedullary hematopoiesis, in which HSCs are mobilized to sites outside the marrow to expand hematopoiesis. Our study showed that the extramedullary splenic hematopoiesis could also compensate/maintain MPN neoplastic hematopoiesis during disease transformation/progression. Together with a previous report that the spleens of PMF patients contain the neoplastic stem cells for MPN development^48^, these findings indicate that effective targeting of the splenic neoplastic hematopoiesis might be necessary for successful MPN therapies.

ECs are an essential component of the perivascular niche in the marrow and spleen^19,45^. It is known that vascular ECs within different tissues have unique gene expression profile and cellular function^49^. Results from our previous studies and current work demonstrated that there was a significant heterogeneity of the JAK2V617F mutant ECs in different parts of the circulation (e.g., marrow, spleen, heart^32^). How the same JAK2V617F mutation results in different EC functions in different tissues is not fully understood. Since flow shear stress has key roles in endothelial function^50^ and biomechanical forces can regulate HSC function^51^, it is possible that different e.g. flow rate, shear stress, or hydrostatic pressure in different tissues can contribute to different JAK2V617F-bearing EC functions.

In summary, our previous^27–32^ and current work have demonstrated that endothelial dysfunction plays an important role in both the hematologic and cardiovascular disease processes of MPNs. The Tie2FF1 mice provide a unique *in vivo* model to screen or test potential preventive and therapeutic interventions for patients with MPNs.

## ACKNOWLEDGEMENTS

This research was supported by the National Heart, Lung, and Blood Institute grant NIH R01 HL134970 (H.Z.) and VA Merit Award BX003947 (H.Z.).

## AUTHOR CONTRIBUTIONS

H. Zhang performed various *in vitro* and *in vivo* experiments of the project, and analyzed the data; A.Y. assisted various *in vitro* culture experiments; S.L. provided technical assistance in various parts of the project; H. Zhan conceived the projects, analyzed the data, interpreted the results, and wrote the manuscript.

